# Molecular architecture of the altered cortical complexity in autism

**DOI:** 10.1101/2024.07.28.605533

**Authors:** Makliya Mamat, Yiyong Chen, Wenwen Shen, Lin Li

## Abstract

Autism Spectrum Disorder (ASD) is characterized by difficulties in social interaction, communication challenges, and repetitive behaviors. Despite extensive research, the molecular mechanisms underlying these neurodevelopmental abnormalities remain elusive. We integrated microscale brain gene expression data with macroscale MRI data from 1829 participants, including individuals with ASD and healthy controls, from the Autism Brain Imaging Data Exchange (ABIDE) I and II. Using fractal dimension (FD) as an index for quantifying cortical complexity, we identified significant regional alterations in ASD, within the left temporoparietal, left peripheral visual, right central visual, left somatomotor (including the insula), and left ventral attention networks. Partial least squares (PLS) regression analysis revealed gene sets associated with these cortical complexity changes, enriched for biological functions related to synaptic transmission, synaptic plasticity, mitochondrial dysfunction, and chromatin organization. Cell-specific analyses, protein-protein interaction (PPI) network analysis and gene temporal expression profiling further elucidated the dynamic molecular landscape associated with these alterations. These findings indicate that ASD-related alterations in cortical complexity are closely linked to specific genetic pathways. The combined analysis of neuroimaging and transcriptomic data enhances our understanding of how genetic factors contribute to brain structural changes in ASD.

## Introduction

Autism Spectrum Disorder (ASD) is a complex neurodevelopmental disorder characterized by a wide range of symptoms, including difficulties in social interaction, communication challenges, and repetitive behaviors (1). The etiology of ASD is multifactorial, involving both genetic and environmental factors (2, 3). Genetic factors are estimated to contribute significantly, between 64% and 93%, to the development of autism (3). Neuroimaging studies have provided valuable insights into the neurobiological basis of ASD, revealing alterations in brain structure and function (4–6). One area of particular interest is cortical changes, as individuals with ASD often exhibit abnormalities in cortical morphology (7). Understanding the molecular changes of the brain in ASD is essential to deciphering the neurobiological underpinnings of ASD, and developing targeted interventions and treatments.

Brain structure can be conceptualized by assessing the intricacy of its surface shape. One of the most promising measures is the cortical complexity, which can be measured through the quantification of fractal dimension (FD) (8, 9). Madan C. R. suggested that fractal dimensionality is the more useful single measure because it simultaneously accounts for shape-related characteristics and serves as a general-purpose measure of structural complexity (10). Cortical complexity, as opposed to measures relying on integral Euclidean geometry, offers a promising method to study the inherent irregularities of cerebral geometry by accounting for the irregular, and fractal convolutedness of cerebral surfaces, thus providing a more suitable approach to capture the natural geometry of the brain (11–14). Notably, cortical complexity analysis offers greater sensitivity in characterizing structural changes compared to conventional volume with small variances and less gender effects (15–17).

Cortical complexity undergoes dynamic changes throughout brain development and aging, increasing during intrauterine and postnatal phases until adolescence and subsequently declining steadily during adulthood (18–21). Hedderich et al. showed decreases in cortical complexity between premature-born adults and full-term controls as a reflection of regionally disturbed neurodevelopmental processes due to premature birth (20). One interesting study revealed that interindividual differences in cortical structure are not only strongly correlated with age, but also can robustly be used to predict age (a combination of cortical thickness and FD showed as the best predictors) (18). In the recent years, cortical complexity measurement demonstrated as a neuroimaging biomarker with high classification accuracy, for brain tumor diagnosis using machine learning (22) and as an input for brain-computer interface (23). Moreover, changes in structural complexity of the cerebral cortex also have been associated with neurological diseases (24–26). Complexity of white matter is also associated with higher fluid cognitive ability (27) and cognitive ability in patients with Alzheimer’s dementia (28).

Few studies have identified altered cortical complexity in individuals with ASD compared to neurotypical controls. Zhao et al. found a significant reduction in the FD of the right cerebellar cortex in ASD relative to typically developing boys (29). Increased cortical complexity was reported in the right fusiform gyrus of the ASD and attention-deficit/hyperactivity disorder (ADHD) cohort in comparison to the ASD-only cohort (30). However, these studies have been limited by small sample sizes and/or quantitative characterization of the entire cortex.

Understanding the structural properties underlying cortical complexity is crucial, as these properties are deeply rooted in genetics. Recent advances in transcriptome imaging analyses have opened new opportunities for understanding how spatial variations on the molecular transcriptomic scale relate to the macroscopic neuroimaging phenotypes by establishing linkages between MRI-based brain measurements and genetic samples obtained from postmortem brains (31, 32). This approach involves mapping gene expression data from the Allen Human Brain Atlas (AHBA) and neuroimaging maps to a common space, such as a parcellated brain atlas (33). Neuroimaging biomarkers are then correlated with the expression levels of thousands of genes in each brain region/network using multivariate statistical methods like partial least squares (PLS) regression (34). Genes are ranked based on how well their expression patterns align with the neuroimaging biomarker under investigation. Researchers have identified dysregulated genes in the post-mortem cortex of individuals with ASD, enriched in synaptic transmission, astrocyte, and microglial functions (35, 36). Romero-Garcia et al. reported a robust association between differences in cortical thickness during childhood and genes involved in synaptic transmission pathways, which are known to be downregulated in the postmortem ASD cortex (37). One recent study identified macroscale changes in cortical networks in autism, further established how macroscale structural connectome alterations in autism relate to microcircuit dysfunction (38). Of most interest for neurodevelopmental disorders is understanding the molecular basis of disorders — to ask, “what causes the differences?” rather than merely “what is different?”

In this context, we sought to bridge gaps by examining alteration of cortical complexity in ASD using large MRI data with 1829 participants, aged between 6.5 and 64 years, from the Autism Brain Imaging Data Exchange (ABIDE) I and II. Moreover, given the tight relationship between cortical structure and gene expression (39), we leveraged brain-wide transcriptomic data from AHBA to identify molecular correlates of ASD-related neuroanatomy irrespective of regional specific neuroanatomical differences. A schematic overview of the study design and analysis pipeline is shown in Figure 1.

**Figure 1.**
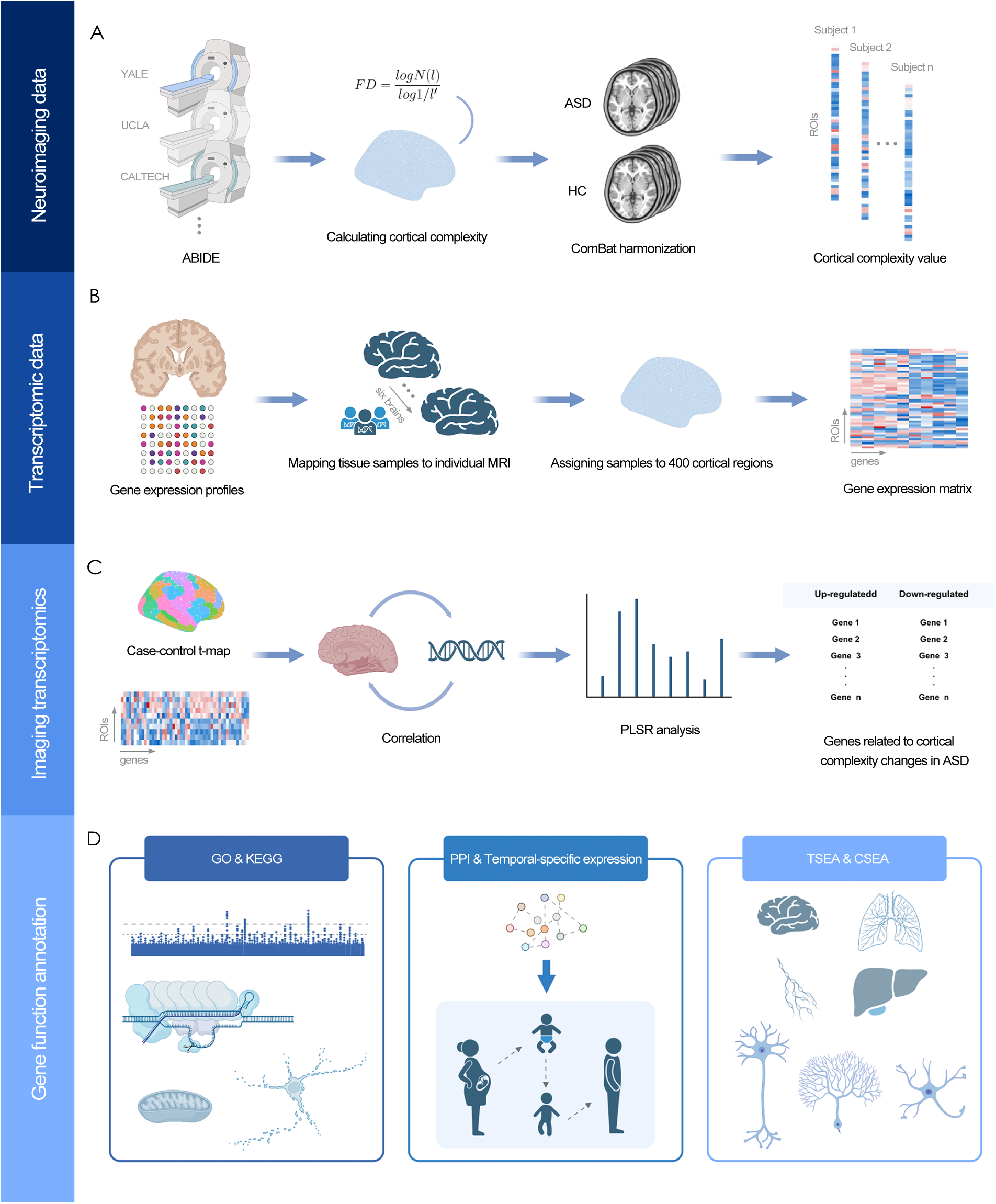
Overview of the Analysis Pipeline. (A) Neuroimaging data processing. Structural MRI data were obtained from the Autism Brain Imaging Data Exchange (ABIDE). Fractal dimension (FD) was computed for each MRI to quantify cortical complexity. To account for site-specific variations, ComBat harmonization was applied. The cortical complexity values across 400 regions of interest (ROIs) were extracted. (B) Transcriptomic Data. Tissue samples were mapped to individual MRIs based on gene expression profiles. These samples were assigned to 400 cortical regions to construct a gene expression matrix, correlating gene activity with specific brain regions. (C) Imaging Transcriptomics Analysis. A case-control t-map was acquired by assessing differences in cortical complexity values between ASD and HC groups. The correlation between cortical complexity and gene expression was assessed using Partial Least Squares Regression (PLSR) analysis. (D) Gene Function Annotation. Gene Ontology (GO) and Kyoto Encyclopedia of Genes and Genomes (KEGG) pathway analyses were performed to elucidate functional pathways. Additionally, protein-protein interaction (PPI) networks and temporal-specific expression patterns were examined.

## Materials and methods

### Participants

The structural MRI data used in this study were obtained from the ABIDE I and II projects (40, 41). All data collection procedures were approved by the local Institutional Review Board. Subject inclusion criteria were as follows: 1) complete whole-brain coverage, 2) good image quality (see follows), and 3) sites with more than 10 subjects in each group after meeting the above criteria. Finally, a total of 1829 subjects (ABIDE I: 460 patients with ASD and 515 healthy controls (HCs) from 15 sites and ABIDE II: 379 patients with ASD and 475 HCs from 13 sites) were included in our study.

Image quality control included two steps: 1) each image was visually inspected for obvious artifacts due to head motion; 2) check the quality control report generated by the Computational Anatomy Toolbox (CAT12) manually, images were excluded if their weighted average image quality rating (IQR) was lower than 70 and volumes with low mean homogeneity (below two standard deviations from the sample mean) were again visually inspected for artefacts. Additional information about the subjects for each site is provided in Table 1. Further information on data acquisition and site-specific details (i.e., protocols, test batteries used, and scanning parameters) is available at the ABIDE website (https://fcon_1000.projects.nitrc.org/indi/abide/).

**Table 1.** Distribution of Participants Across Different Sites in ABIDE I and II Datasets. ABIDE, Autism Brain Imaging Data Exchange; ASD, autism spectrum disorder; HC, healthy control; BNI, Barrow Neurological Institute; CALTECH, California Institute of Technology; CMU, Carnegie Mellon University; EMC, Erasmus University Medical Center Rotterdam; GU, Georgetown University; IP, Institut Pasteur and Robert Debré Hospital; IU, Indiana University; KKI, Kennedy Krieger Institute; LEUVEN, University of Leuven; MAX_MUN, Ludwig Maximilians University Munich; NYU, New York University Langone Medical Center; OHSU, Oregon Health and Science University; OLIN, Institute of Living at Hartford Hospital; PITT, University of Pittsburgh School of Medicine; SBL, Social Brain Lab BCN NIC UMC Groningen and Netherlands Institute for Neurosciences; SDSU, San Diego State University; SU, Stanford University; TCD, Trinity Centre for Health Sciences; TRINITY, Trinity Centre for Health Sciences; UCD, University of California Davis; UCLA, University of California, Los Angeles; UM, University of Michigan; USM, University of Utah School of Medicine; YALE, Yale Child Study Center, Yale School of Medicine.

| ABIDE I |  |  | ABIDE II |  |  |
| --- | --- | --- | --- | --- | --- |
| Site | ASD | HC | Site | ASD | HC |
| CALTECH | 18 | 19 | BNI | 29 | 29 |
| CMU | 14 | 13 | EMC | 20 | 22 |
| KKI | 20 | 32 | GU | 40 | 49 |
| LEUVEN | 29 | 33 | IP | 15 | 30 |
| MAX_MUN | 22 | 32 | IU | 18 | 20 |
| NYU | 75 | 102 | KKI | 52 | 150 |
| OHSU | 13 | 15 | NYU | 66 | 27 |
| OLIN | 16 | 15 | OHSU | 37 | 55 |
| PITT | 29 | 27 | SDSU | 32 | 25 |
| SBL | 14 | 14 | SU | 21 | 21 |
| TRINITY | 23 | 25 | TCD | 18 | 20 |
| UCLA | 50 | 42 | UCD | 14 | 12 |
| UM | 53 | 75 | USM | 17 | 15 |
| USM | 57 | 43 |  |  |  |
| YALE | 27 | 28 |  |  |  |
| Total | 460 | 515 | Total | 379 | 475 |

### Magnetic resonance imaging data pre-processing

T1 images were manually set the origin at the anterior commissure, then were processed using the CAT12 toolbox (version 1980, Structural Brain Mapping, Jena University Hospital, Jena, Germany) implemented in SPM12 (version 7771, Institute of Neurology, London, UK). We employed the default parameters of CAT12 for this pre-processing procedure. All the T1-weighted images were corrected for bias-field inhomogeneities, then segmented into gray matter, white matter, and cerebrospinal fluid and spatially normalized using the DARTEL algorithm (42). The final resulting voxel size was 1.5 × 1.5 × 1.5 mm.

For surface-based morphometry, CAT12 toolbox computes multiple surface parameters, including FD (43). These surface parameters of the left and right hemispheres were separately resampled and smoothed with a 20-mm FWHM Gaussian kernel. The software parcellated the cortex into 400 regions of interest (ROI) using the Schaefer atlas 17 networks for surface measures (44). We then averaged the FD value from each ROI in each participant for further analyses.

### Mega analysis

As the ABIDE datasets are multicentric with heterogeneous acquisition parameters across sites, raw FD values were harmonized between sites using the ComBat method to remove inter-site variation and preserve biological variability (45). Independent two-sample t-tests were performed between ASD and HC groups to identify cortical complexity changes related differences. All statistical analyses were performed using R software (version 4.2, https://www.r-project.org/), and a threshold of p < 0.05, False Discovery Rate (FDR) corrected, was applied.

### Meta analysis

To account for differences in scanners, acquisitions and sample characteristics, statistical analysis was conducted using a prospective meta-analytic technique, where each site is initially treated as an independent study and results are pooled to define significance. Effect sizes were computed as standardized mean differences (Cohen’s d) using Hedges’g as estimator. The between-study variance τ^2^ was estimated using the restricted maximum likelihood method, from which we computed the proportion of variance imputable to heterogeneity. Computations were performed using R (https://www.r-project.org) with the packages meta and metafor (46). We report statistical significance for an α level of.05.

### Gene expression data processing

Gene expression data were obtained from the AHBA (http://human.brain-map.org). The AHBA comprises the normalized expression data of 20737 genes represented by 58692 probes taken from 3702 brain tissue samples from six donors (one female and five males, aged 24–57 years) (32). A newly proposed pipeline for transcription-neuroimaging association studies based on AHBA data was used in this study (33). Only genes that were consistently expressed across donors (i.e., average inter-donor correlation ≥0.5) were considered for our analyses. To correct for donor-specific effects, scaled robust sigmoid (SRS) normalization was used to ensure equivalent scaling of expression values for each donor. After this procedure, the expression values were more comparable across donors. Finally, we obtained a normalized gene expression matrix of 400 × 15633 (ROI × gene) (44). The detailed preprocessing steps are described in Supplementary Material.

### Identifying transcriptomic correlates of cortical complexity changes in ASD

To identify genes whose expression was inferentially correlated with ASD-related alterations, we used a PLS regression, a multi-variate technique accounting for inherent shared topological structure between brain-derived neuroimaging phenotypes and gene expression. Significant genes were obtained by regressing each gene against our t-statistical maps and using a one-sample t-test to determine whether the slopes were different from 0. To correct for multiple comparisons, the procedure was repeated by randomly rotating our maps using 1000 spin permutations, which were compared with the original t-statistic to assess gene significance. We then ranked all genes according to their z score weights to the PLS components.

From the ranked PLS gene list, genes with Z score more than 1.96 or less than −1.96 (*P_FDR_* < 0.05, FDR corrected for the total number of genes tested), were selected. These are denoted as PLS+ and PLS-, representing genes most positively and negatively associated with FD changes in ASD patients.

### Gene enrichment analyses

We employed the Metascape software to perform Gene Ontology (GO) and Kyoto Encyclopedia of Genes and Genomes (KEGG) Pathways analysis (https://metascape.org/) (47). GO was used to determine their biological functions including molecular functions (MFs), biological processes (BPs), and cellular components (CCs) (48). KEGG was used to identify related biological pathways (49).

We employed Specific Expression Analysis (SEA) (50) to assess the potential over-representation of cortical complexity changes related genes in three specific domains, namely cell types, brain regions, and developmental stages. This analysis incorporated a specificity index probability (pSI), offering insights into the enrichment levels of genes within specific terms compared to others (51). The transcriptional profiles for brain development were sourced from the BrainSpan Atlas of the Developing Human Brain (http://www.brainspan.org/), spanning 13 developmental stages across 8–16 brain structures. All enrichment analyses were executed using Fisher’s exact tests, and Benjamini-Hochberg False Discovery Rate (BH-FDR) correction was applied to account for multiple comparisons, ensuring a stringent threshold of significance (q < 0.05) for gene functional annotations.

We used TissueEnrich R package (52) and cell type specific expression analyses (CSEA) tools (http://doughertytools.wustl.edu/CSEAtool.html) (51) to conduct tissue, cell type, and temporal specific expression analyses. These specific expression analyses could help to determine the specific tissues, cortical cell types, and developmental stages in which the FD alteration related genes were overrepresented. Fisher’s exact tests were used to assess the significance of the above-mentioned enrichment analyses. Multiple testing was corrected using the BH-FDR correction with a corrected P value of 0.05.

We constructed PPI networks form the up and down regulated gene sets using STRING version 10.5, with the highest confidence value of 0.9 (53). Hub genes were defined by the top 1% of the node degree in the PPI networks. Additionally, the Human Brain Transcriptome database (http://hbatlas.org/) was used to characterize the spatial–temporal expression trajectory of hub genes with the highest node degree.

## Results

### Case-control differences

Mega analysis revealed significant alterations (P < 0.05, FDR corrected) in cortical complexity between individuals with ASD and HC in several key brain networks (Fig. 2A). The case-control difference pattern from the meta-analysis (Supplementary Fig. 1) was remarkably similar to that derived from the mega-analysis (spatial similarity: *r* = 0.95, *p* <.0001) (Fig. 2B). Notable differences were observed in the left temporoparietal network, and in the left peripheral visual network. Additionally, significant changes were found in the right central visual network, the left somatomotor network, particularly within the insula, and the left ventral attention network (Fig. 2C).

**Figure 2.**
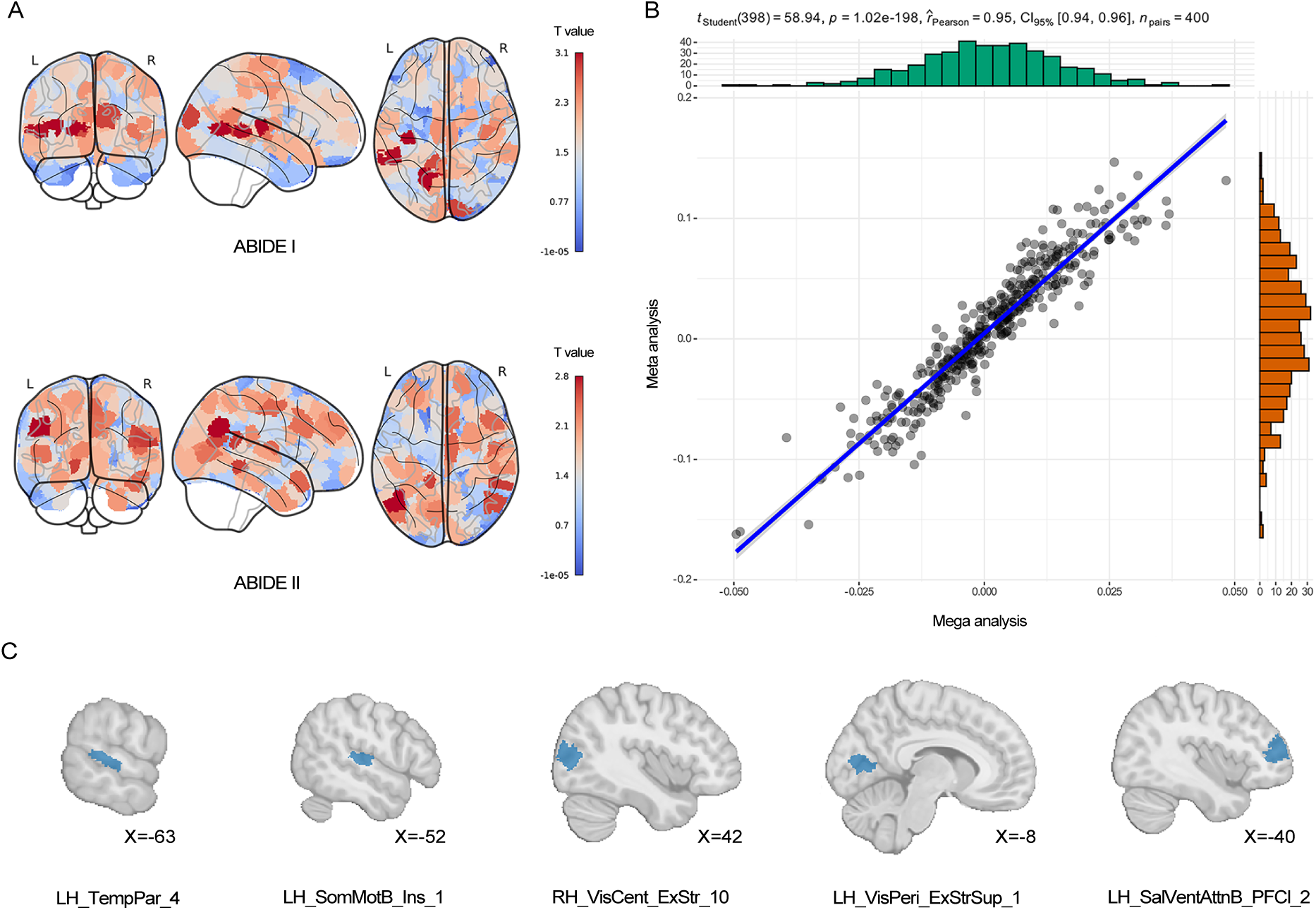
Case-Control Differences in Cortical Complexity. (A) T-maps of cortical complexity differences. T-maps for ABIDE I and ABIDE II datasets highlight regions with significant cortical complexity differences between autism spectrum disorder (ASD) and healthy control (HC) groups. (B) Spatial similarity between case-control difference maps obtained from the mega-and meta-analyses. (C) The Schaefer brain networks showing significant cortical complexity alterations in ASD compared to HC are highlighted in blue. ABIDE, Autism Brain Imaging Data Exchange; LH, left hemisphere; RH, right hemisphere; TempPar, temporoparietal network; SomMotB_Ins, somatomotor network including insula; VisCent_ExStr, central visual network; VisPeri_ExStrSup, peripheral visual network; SalVentAttnB_PFCi, ventral attention network.

### Brain gene expression profiles associated with cortical complexity changes

We investigated the relationship between brain-wide gene expression maps and cortical complexity changes in ASD using a PLS regression analysis. The first and third PLS components of ABIDE I and the first component of ABIDE II were extracted based on their high statistical significance (P < 0.05), embodying linear combinations of weighted gene expression scores associated with the t-statistical map (Fig. 3, Dataset 1 listed the full gene terms before the Z score cut-off). Subsequently, upregulated and downregulated gene sets were meticulously extracted from the first and third components of ABIDE I, along with corresponding gene sets from the first component of ABIDE II. Intersections of upregulated and downregulated genes were determined, resulting consolidated lists capturing genes associated with up regulated genes (denoted as URGs) and down regulated genes (denoted as DRGs) related to cortical complexity changes in ASD, forming the basis for subsequent enrichment analyses.

**Figure 3.**
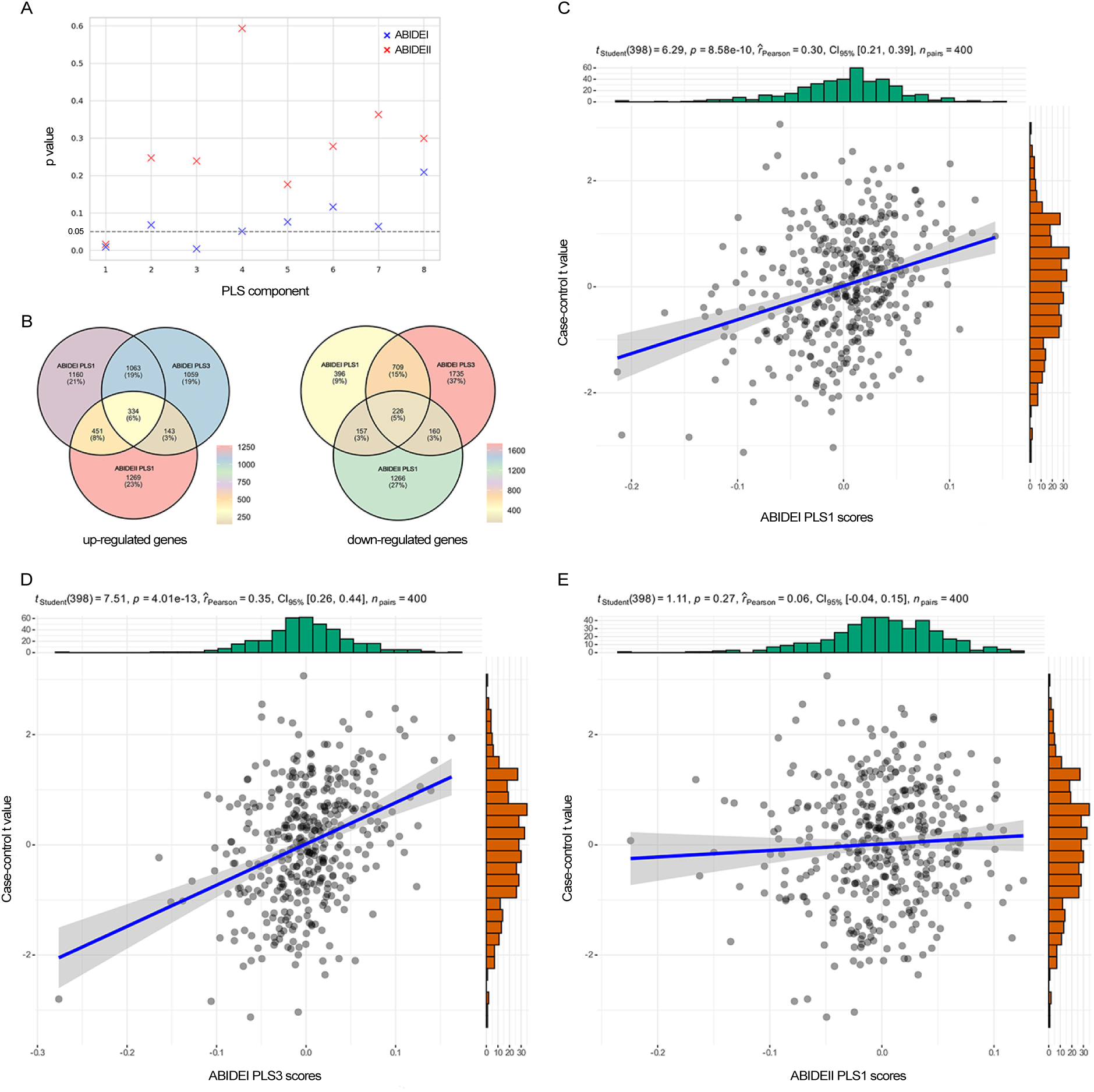
Differential Gene Expression Analyses. (A) PLS Component Analysis. The plot depicts the p-values of PLS components for ABIDE I (blue) and ABIDE II (red) datasets, indicating the statistical significance of each component. (B) Venn Diagrams of Up-and Down-Regulated Genes. The left shows the overlap of up-regulated genes between ABIDE I and ABIDE II datasets across different PLS components, while the right illustrates the overlap of down-regulated genes between the two datasets. (C) The plot shows the correlation between case-control values and PLS1 scores for ABIDE I. (D) The plot shows the correlation between case-control values and PLS3 scores for ABIDE I. (E) The plot depicts the correlation between case-control values and PLS1 scores for ABIDE II. ABIDE, Autism Brain Imaging Data Exchange; PLS, partial least squares.

### Gene enrichment analyses

We specifically focused on aligning functional enrichments with URGs. Our analysis revealed significant enrichment of URGs in several key biological processes, including modulation of synaptic transmission, signaling, and nervous system processes. Additionally, URGs exhibited enrichment in cellular components such as the mitochondrial membrane and respirasome, and in molecular functions including electron transfer activity and inorganic molecular entity transmembrane transporter activity. Meanwhile, DRGs showed significant enrichment in biological processes associated with chromatin organization and regulation of cellular response to stress. Furthermore, DRGs exhibited enrichment in cellular components such as the centrosome and Golgi membrane, and in molecular functions including histone binding and molecular adaptor activity. Notably, our analysis also identified an association of DRGs with neurodegeneration pathways, as revealed by KEGG analysis. Figure 4A-B showed representative gene enrichment terms for URGs (see Dataset 2 for full list).

**Figure 4.**
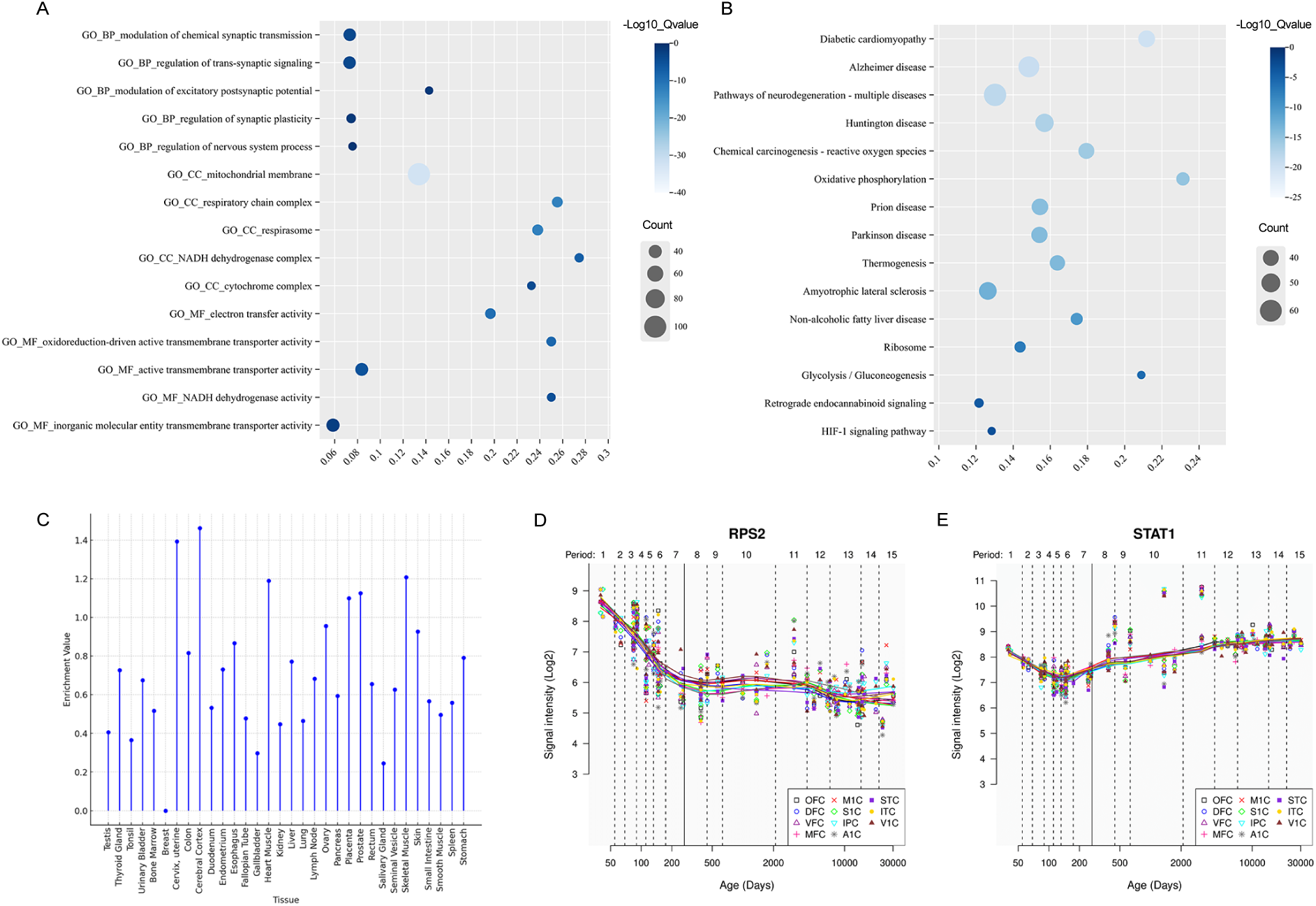
Functional Enrichment and Expression Analysis. (A) Dot plot shows enriched Gene Ontology (GO) terms of up regulated genes (URGs). (B) Dot plot illustrates enriched Kyoto Encyclopedia of Genes and Genomes (KEGG) pathways of URGs. (C) Lollipop chart indicates the tissue-specific expression patterns of the identified URGs. (D) Line plots show the temporal expression patterns of RPS2 across different developmental stages. (E) Line plots show the temporal expression patterns of STAT1 across different developmental stages. BP, biological process; CC, cellular components; MF, molecular function; OFC, orbital prefrontal cortex; DFC, dorsolateral prefrontal cortex; VFC, ventrolateral prefrontal cortex; MFC, medial prefrontal cortex; M1C, primary motor (M1) cortex; S1C, primary somatosensory (S1) cortex; IPC, posterior inferior parietal cortex; A1C, primary auditory (A1) cortex; STC, superior temporal cortex; ITC, inferior temporal cortex; V1C, primary visual (V1) cortex.

### Tissue and cell type specific expression

The URGs and DRGs related to cortical complexity changes in ASD exhibited specific expression patterns in brain tissue, including in the cerebral cortex (Fig. 4C and Supplementary Fig. 2, 3). SEA of adult brain regions demonstrated significant expression of the identified gene sets in key brain regions, including the cerebellum, cerebral cortex, thalamus, and hippocampus. Moreover, cell type SEA revealed higher expression levels in specific neuronal populations, such as Pnoc+ neurons in the cortex and cholinergic neurons in the striatum (Supplementary Fig. 4, 5). Additionally, developmental gene expression analysis indicated that URGs were expressed in the brain from late fetal development onward, across various brain regions including the cortex, and subcortex (hippocampus, striatum, thalamus), whereas DRGs exhibited expression from early to mid-fetal development onward, across several brain regions including the cortex and subcortex (amygdala, thalamus).

### PPI networks, hub genes, and temporal specific expression

We constructed networks of known interactions between proteins encoded by the two gene sets (Supplementary Fig. 6, 7). For the URGs, the resulting network comprised 889 nodes and 966 edges, significantly more than the 654 edges expected by chance, with an average node degree of 2.17 (PPI enrichment P-value < 1.0e-16). Similarly, for the DRGs, the resulting network consisted of 492 nodes and 93 edges, surpassing the expected 73 edges, with an average node degree of 0.378 (PPI enrichment P-value 0.0151). For detailed information on the constructed network see Dataset 3.

Hub genes were identified as those within the top 1% of the node degree in each PPI network. In total, we identified 9 and 5 hub genes involved in the PPI networks constructed by the URGs and DRGs, respectively (Supplementary Fig. 8). Here, we characterized the spatial-temporal expression trajectory of two hub genes with the highest node degree for each gene set, namely RPS2 for the URGs and STAT1 for the DRGs (Fig. 4).

## Discussion

We explored the molecular underpinnings of cortical complexity alteration by integrating microscale brain gene expression with macroscale MRI data from multi-scanner large datasets. We identified significant alterations in cortical complexity between individuals with ASD and HC in several key brain regions, including the left temporoparietal network, left peripheral visual network, right central visual network, left somatomotor network (particularly within the insula), and left ventral attention network. The first and third PLS components of ABIDE I and the first component of ABIDE II explain a significant proportion of variance in these cortical complexity alterations. We extracted two gene sets that positively (URGs) and negatively (DRGs) associated with the alteration of cortical complexity in ASD relative to the healthy controls. These genes were significantly enriched for biological functions and pathways related to synaptic transmission, synaptic plasticity, mitochondrial dysfunction, neurodegeneration pathways, electron transfer activity, and chromatin organization and remodeling. Specific expression analyses revealed that these cortical complexity changes related genes were expressed in brain tissue, particularly in cortical neurons, across various developmental periods. Additionally, PPI analysis revealed that these genes could construct a PPI network with 14 hub genes.

In this study, we employed FD as an index for quantifying cortical complexity. It has been shown to be a favorable complement to other analyses of cortical complexity alterations, with greater sensitivity (54). Fractal dimensionality appears to be a highly reliable structural measure (55), and capable of differentiating individuals with various psychiatric and neurological disorders from healthy controls (17, 28, 56). Several significant alterations in cortical complexity between individuals with ASD and HCs have been identified in our study. The left temporoparietal network, which is crucial for integrating sensory and cognitive information, showed significant alterations in individuals with ASD. This network is involved in processes such as language comprehension, social cognition, and theory of mind (57). The observed differences in this region may underlie some of the core deficits in social interaction and communication seen in ASD. Previous research has indicated abnormal functional connectivity in the temporoparietal junction in ASD, which is associated with difficulties in understanding others’ perspectives and social cues (58, 59). Significant changes were also found in both the left peripheral visual network and the right central visual network. The peripheral visual network is essential for higher-order visual processing, including the integration of complex visual stimuli, while the central visual network is critical for primary visual processing (60, 61). Alterations in these networks suggest potential disruptions in visual processing pathways in ASD, which may contribute to the atypical experiences and perceptual processing often reported in individuals with ASD (62, 63).

The left somatomotor network, particularly within the insula, exhibited significant changes. The insula plays a vital role in sensorimotor integration and emotional processing (64). Alterations in this region may be linked to the sensory processing abnormalities and emotional dysregulation frequently observed in ASD (65). This finding aligns with previous studies that have reported atypical insular activity and connectivity in ASD, associated with sensory overresponsivity and difficulties in emotional awareness (66). The left ventral attention network, involved in executive functions and attentional control, also showed significant alterations. This network is crucial for detecting and responding to salient stimuli in the environment (67). Changes in this network may contribute to the executive function deficits and attentional dysregulation observed in ASD (68, 69). Previous studies have demonstrated that individuals with ASD often exhibit difficulties in shifting attention and managing distractions, which are linked to abnormalities in the ventral attention network (70, 71).

The observed alterations in specific brain networks suggest potential targets for intervention and therapy. Future research should aim to replicate these findings in larger, independent samples and explore the longitudinal trajectory of cortical complexity changes across different developmental stages in ASD. Additionally, integrating multimodal imaging techniques and genetic analyses could provide a more comprehensive understanding of the neurobiological and molecular mechanisms underlying ASD. Understanding how these structural changes relate to functional outcomes and behavioral phenotypes will be crucial in developing personalized and effective interventions for individuals with ASD.

One plausible explanation for the observed cortical complexity alterations in multiple brain regions in ASD compared to healthy controls is aberrant neurodevelopment. It has been suggested that during brain development, individuals with ASD may experience disruptions in neuronal proliferation, migration, and cortical organization, leading to atypical cortical complexity patterns (72). Studies have reported abnormalities in gene expression, neuronal connectivity, and synaptic pruning in ASD, which could contribute to alterations in cortical morphology (35, 73).

Another potential explanation is altered neuronal connectivity in individuals with ASD. Diffusion tensor imaging studies have consistently reported disruptions in white matter tracts and connectivity patterns in ASD (74). The disruption of white matter connectivity may directly account for the cortical complexity changes, given that tension along axons in the white matter is the primary driving force for cortical folding (75). Interestingly, alterations in grey matter cortical complexity have been suggested to be secondary to axonal damage in the white matter (56). Zhao et al. reported that male children with ASD exhibit significantly reduced structural complexity in the right cerebellar cortex compared to healthy controls (29). These alterations in connectivity could affect the development and organization of cortical networks, thereby influencing cortical complexity (76). Nicastro and his collogues proposed that changes in cortical complexity may depend on the extent of structural impairment affecting the pial surface. A decrease in complexity is more likely if alterations in the pial surface reduce the folding area, while an increase in complexity would be expected if the change involves an increase in sulcal depth (77). Moreover, both environmental and genetic factors could contribute to the observed alterations in cortical complexity. Prenatal and perinatal environmental factors, such as maternal immune activation and exposure to toxins, have been implicated in the etiology of ASD and could influence cortical development (78). Numerous genetic studies have identified risk genes associated with ASD, many of which are involved in neurodevelopmental processes and synaptic function (79, 80).

Given the heterogeneity in autism imaging findings (2), it is pertinent to ask how genetic risk for autism is associated with variability in cortical complexity changes. Two distinct studies offered converging evidence indicating that alterations in both cortical thickness (37) and regional volumes (81) are associated with the expression of genes that are downregulated in postmortem cortical tissue of individuals with ASD. The regional distribution of cortical thickness alterations observed in ASD in their study was also associated with synaptic transmission genes, whereas changes in regional volumes were associated with nervous system and ion transport–related genes.

In our study, the enrichment of URGs involved in chemical synaptic transmission and excitatory postsynaptic potential aligns with previous research implicating synaptic dysregulation in ASD pathology (35, 37). Synaptic processes play a critical role in ASD, where aberrant synaptic function is associated with impaired communication between neurons and disrupted network dynamics (82). Numerous candidate genes linked to ASD are involved in synaptic architecture and function, mediating the transmission of information between neurons and other cell types (83). Previous investigations have identified disruptions in trans-synaptic signaling as a common theme in ASD, potentially impacting information processing and connectivity within neural networks (84). Synaptic plasticity, a fundamental aspect of learning and memory, has been extensively studied in the context of ASD, with evidence suggesting that aberrations in plasticity mechanisms contribute to cognitive and behavioral phenotypes associated with the disorder (85, 86). The collective enrichment of synaptic terms underscores the potential regulatory roles of the identified genes in governing synaptic dynamics, thereby contributing to the observed alterations in cortical complexity in individuals with ASD.

Researchers have established the involvement of a wide array of genes in nervous system processes associated with neurodevelopmental disorders (87, 88). Furthermore, our study revealed the enrichment of genes associated with mitochondrial components, such as the mitochondrial membrane, respiratory chain complex, and respirasome, suggesting a potential link between altered cortical complexity and mitochondrial functions in ASD. There is a growing body of evidence highlighting mitochondrial dysfunction as a contributing factor in the pathophysiology of ASD (89). Mitochondrial dysfunction may impact cellular energetics, influencing processes essential for proper neurodevelopment. This association is particularly relevant, given the crucial role of mitochondria in synaptic transmission, neuronal plasticity, and overall neuronal health (90).

Neurodegeneration pathways encompass a wide array of cellular processes, including protein aggregation, oxidative stress, mitochondrial dysfunction, and synaptic dysfunction, among others. The identification of neurodegeneration pathways enriched among URGs associated with cortical changes in ASD suggests potential shared molecular pathways between ASD and neurodegenerative disorders such as Alzheimer’s disease, Parkinson’s disease, and Huntington’s disease. This shared molecular landscape may offer insights into common underlying mechanisms contributing to altered brain structure and function across different neurodevelopmental and neurodegenerative conditions. The identification of GO terms associated with electron transfer activity, particularly involving the NADH dehydrogenase complex, and oxidoreduction-driven active transmembrane transporter activity within our transcriptomic analysis hints at putative implications for cellular energetics and redox homeostasis (91, 92). These observations show a plausible role for the identified genes in modulating fundamental metabolic pathways crucial for sustaining cellular function and homeostasis within the cerebral cortex.

We observed significant enrichment of DRGs in chromatin organization and remodeling, processes critical for regulating gene expression patterns during neurodevelopment and essential for proper neuronal differentiation and maturation. (93, 94). Additionally, chromosome segregation defects and impaired stress response mechanisms have been implicated in various neurodevelopmental disorders, including ASD (95, 96). The enrichment of cellular components such as the centrosome, Golgi membrane and endosome membrane suggests disruptions in intracellular trafficking and organelle dynamics in ASD. These cellular components are integral to various cellular processes, including protein trafficking, organelle biogenesis, and cell division, all of which are crucial for proper neuronal function and brain development (97, 98). Dysregulation of these molecular functions can impact gene expression profiles and contribute to the observed changes in neuronal connectivity and synaptic function associated with cortical complexity changes in ASD.

Our study has some limitations worth noting. First, although we combined the large datasets ABIDE I and II and identified PLS components with significant p-values, studies using different validations are needed. Additionally, the challenge of characterizing the specificity of identified associations. The spatial expression pattern of certain gene categories may lead to them being identified as significantly enriched. However, the limited number of specimens in the AHBA dataset prevents a detailed study of individual differences in gene expression. Although imaging transcriptomics can initially identify molecular correlates of anatomically varying cortical complexity, further investigation is required to understand how these correlates vary among individual brains (99). Furthermore, the whole-brain gene expression data utilized in this study were derived from only six postmortem adult brains, with data from the right hemisphere obtained from only two donors. Despite our focus on probes selected to optimize differential stability across donors, the limited sample size restricts the extent to which strong assertions can be made regarding the stability of gene expression across human brains.

In summary, we identified cortical complexity alterations in ASD compared to healthy controls and explored the underlying genetic determinants. We identified genes and transcript expression changes in ASD that occur across the cerebral cortex, affecting many neural cell types and specific biological processes. As we seek to gain a deeper understanding of cortical complexity changes in ASD, future approaches that integrate different sources of biological data and more specific methods to determine how ASD risk genes affect the brain structure will be essential.

## Supporting information

Data set 1

Data set 2

Data set 3

## Declaration of interests

The authors declare no competing interests.

## Acknowledgments

We would like to thank the Autism Brain Imaging Data Exchange (ABIDE) consortium for providing the invaluable dataset that made this research possible. Their efforts in compiling and maintaining the dataset have greatly facilitated advancements in autism research. This work was supported by the Municipal Key R&D Program of Ningbo (2023Z175), and the Applied Basic Research Programs of Science and Technology Commission Foundation of Zhejiang Province (BD23H090003).

## Notes

### Competing Interest Statement

The authors have declared no competing interest.

### Summary of Updates

Figure 1 revised for better image resolution

