## Supplementary Material for "Molecular architecture of the altered cortical complexity in autism"

### Gene Expression Data Processing

Regional microarray expression data were obtained from 6 post-mortem brains (1 female, ages 24.0–57.0, 42.50 +/- 13.38) provided by the Allen Human Brain Atlas (AHBA, https://human.brain-map.org; (1)). Data were processed with the abagen toolbox (version X.Y; https://github.com/rmarkello/abagen) using a 400-region volumetric atlas in MNI space (2).

First, microarray probes were reannotated using data provided by (3); probes not matched to a valid Entrez ID were discarded. Next, probes were filtered based on their expression intensity relative to background noise (4), such that probes with intensity less than the background in >=50.00% of samples across donors were discarded. When multiple probes indexed the expression of the same gene, we selected and used the probe with the most consistent pattern of regional variation across donors (i.e., differential stability; (5)), calculated with:

$$\Delta_S(p) = \frac{1}{\binom{N}{2}} \sum_{i=1}^{N-1} \sum_{j=i+1}^{N} \rho\left[B_i(p), B_j(p)\right]$$

where $\rho$ is Spearman's rank correlation of the expression of a single probe, p, across regions in two donor brains $B_i$ and $B_j$, and N is the total number of donors. Here, regions correspond to the structural designations provided in the ontology from the AHBA.

The MNI coordinates of tissue samples were updated to those generated via non-linear registration using the Advanced Normalization Tools (ANTs; https://github.com/chrisfilo/alleninf). Samples were assigned to brain regions in the provided atlas if their MNI coordinates were within 2 mm of a given parcel. To reduce the potential for misassignment, sample-to-region matching was constrained by hemisphere and gross structural divisions (i.e., cortex, subcortex/brainstem, and cerebellum, such that e.g., a sample in the left cortex could only be assigned to an atlas parcel in the left cortex; [A2019N]). All tissue samples not assigned to a brain region in the provided atlas were discarded.

Inter-subject variation was addressed by normalizing tissue sample expression values across genes using a robust sigmoid function (6):

$$x_{norm} = \frac{1}{1 + \exp(-\frac{(x - \langle x \rangle))}{IQR_x})}$$

where $\langle x \rangle$ is the median and $IQR_x$ is the normalized interquartile range of the expression of a single tissue sample across genes. Normalized expression values were then rescaled to the unit interval:

$$x_{scaled} = \frac{x_{norm} - \min(x_{norm})}{\max(x_{norm}) - \min(x_{norm})}$$

Gene expression values were then normalized across tissue samples using an identical procedure. Samples assigned to the same brain region were averaged separately for each donor and then across donors.

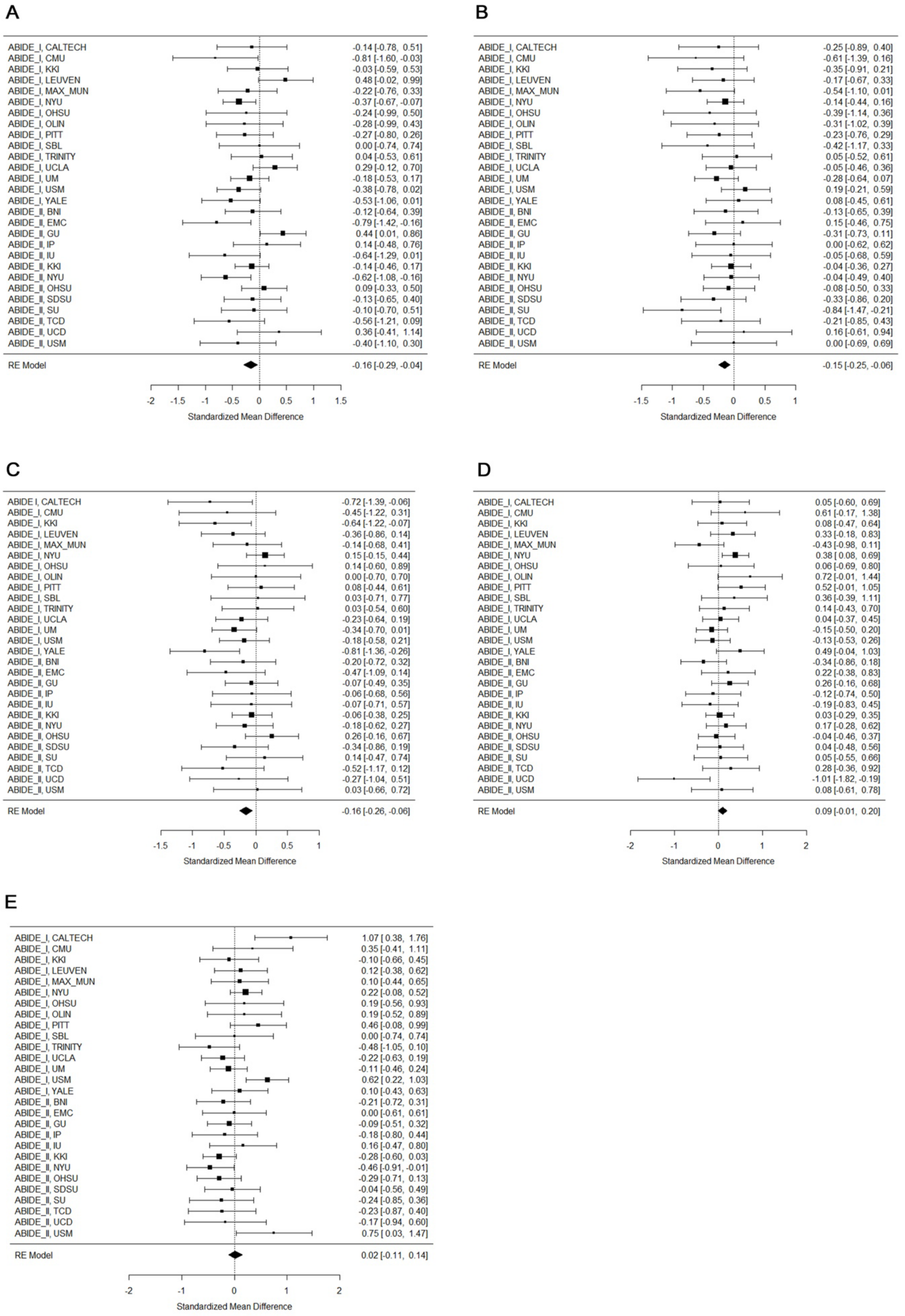


**Supplementary Figure 1.** Meta-analysis of case-control differences in cortical complexity. Forest plot of Cohen's *d* effect sizes for case-control differences in the left peripheral visual network (A), right central visual network (B), left temporoparietal network (C), left somatomotor network (D), left ventral attention network (E).

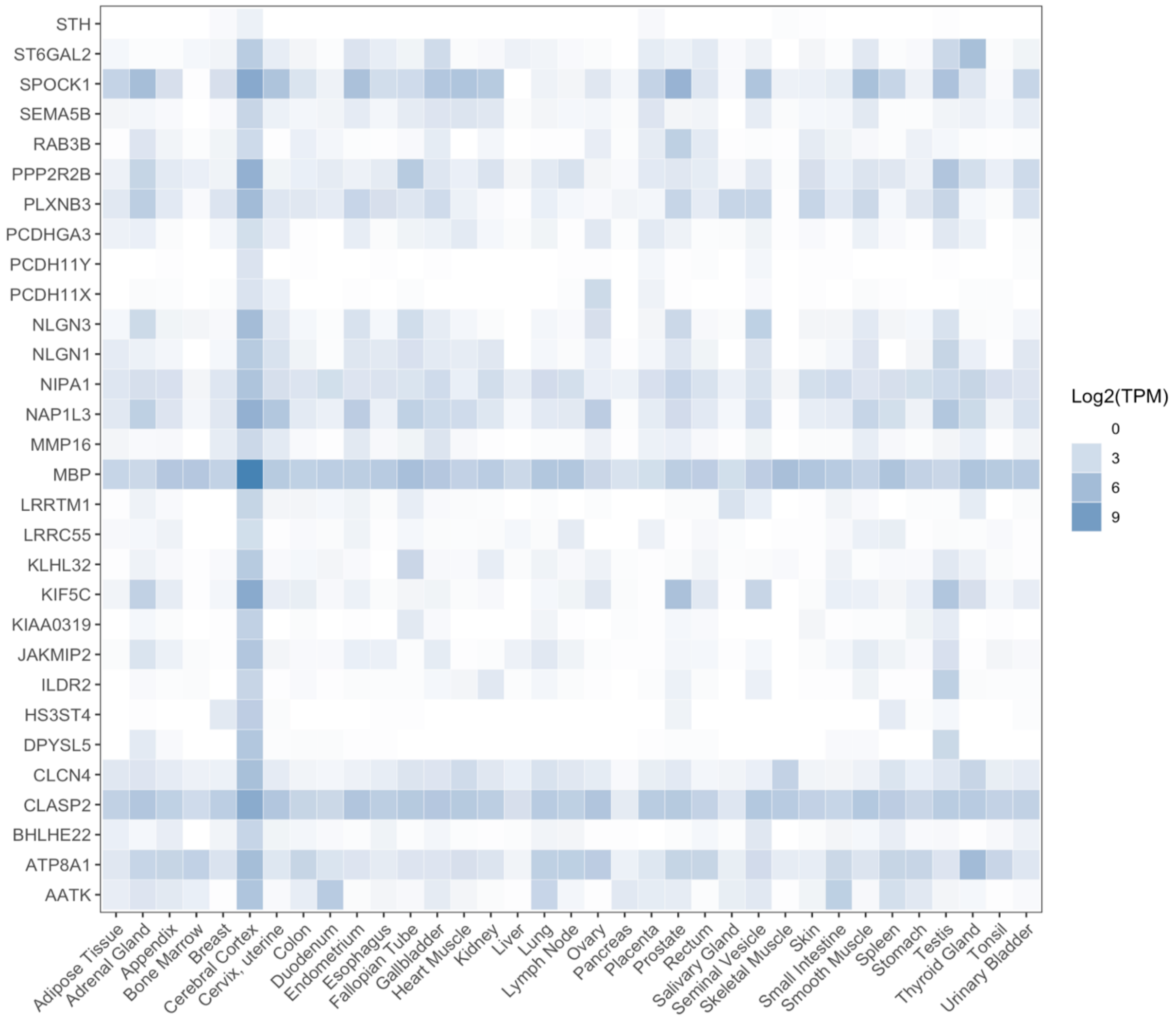


**Supplementary Figure 2.** Heatmap shows the expression profiles of URGs in association with changes in cortical complexity in ASD. TPM, transformed transcripts per million.

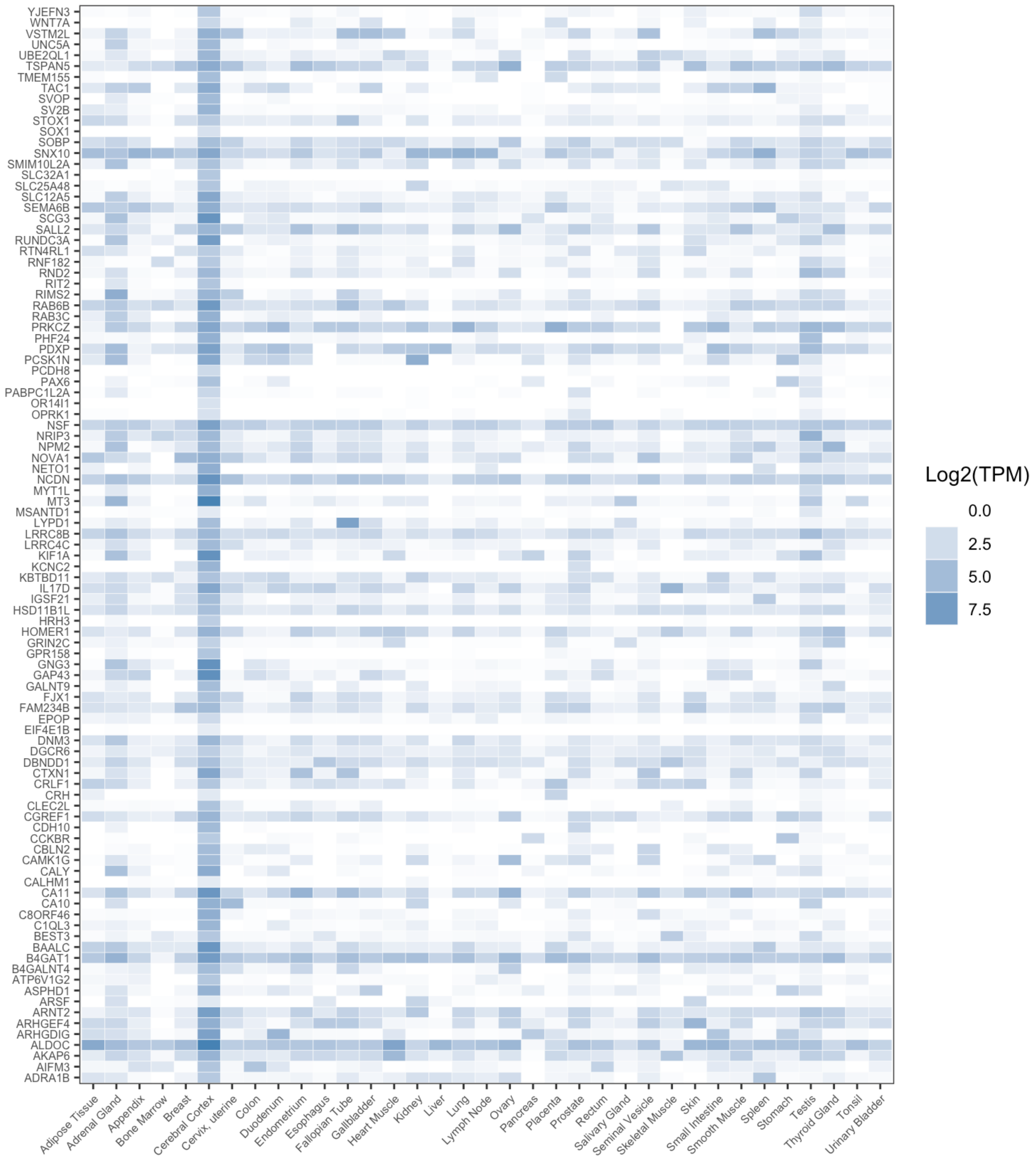


**Supplementary Figure 3.** Heatmap shows the expression profiles of DRGs in association with changes in cortical complexity in ASD. TPM, transformed transcripts per million.

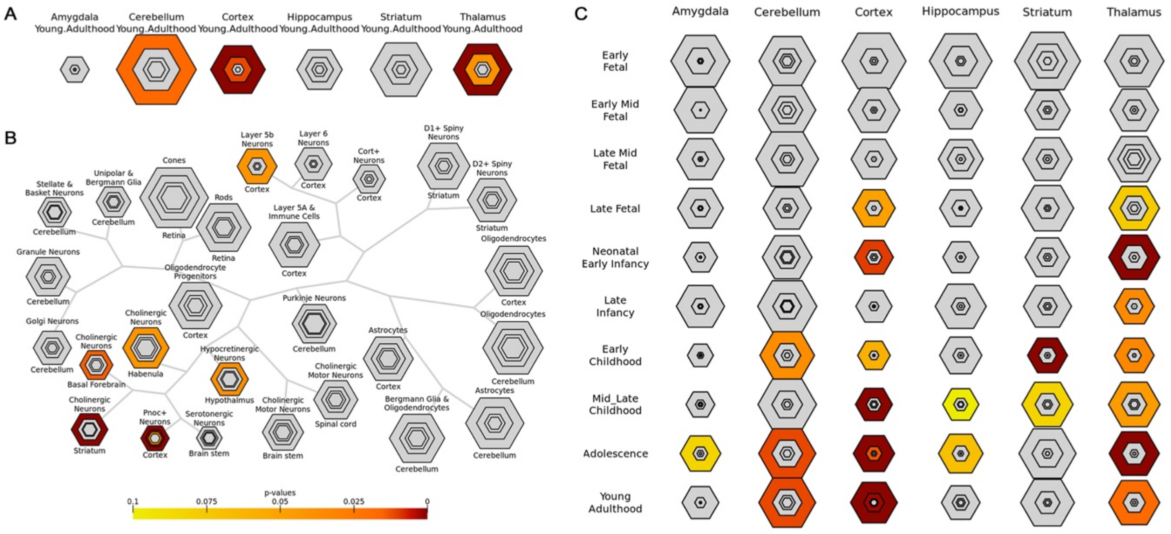


**Supplementary Figure 4.** Specific expression analyses (SEA) of the up-regulated genes related to the cortical complexity changes in ASD. (A) Brain region SEA. (B) Cell type SEA. (C) Development SEA. Colored cell types represent significance after multiple testing correction. The sizes of the hexagons denote cell-type specificity across different specificity index probability (pSI) statistic thresholds ranging from 0.05 to 1e-4. The outer hexagons correspond to the least specific test for a cell type (pSI threshold = 0.05), whereas the innermost hexagon reflects the most specific test for a cell type (pSI threshold = 1e-4).

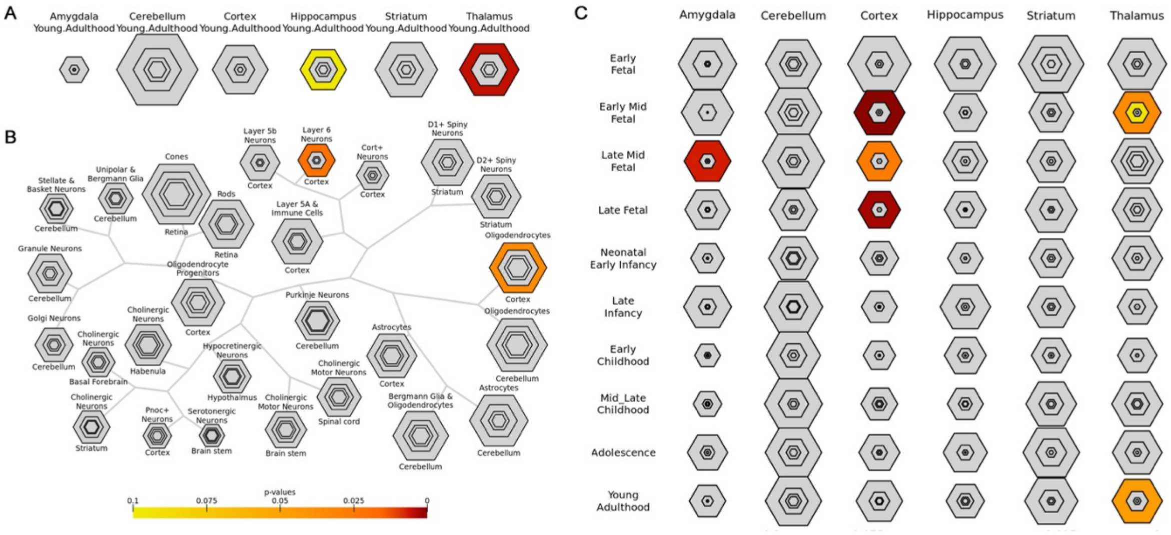


**Supplementary Figure 5.** Specific expression analyses (SEA) of the down-regulated genes related to the cortical complexity changes in ASD. (A) Brain region SEA. (B) Cell type SEA. (C) Development SEA. Colored cell types represent significance after multiple testing correction. The sizes of the hexagons denote cell-type specificity across different specificity index probability (pSI) statistic thresholds ranging from 0.05 to 1e-4. The outer hexagons correspond to the least specific test for a cell type (pSI threshold = 0.05), whereas the innermost hexagon reflects the most specific test for a cell type (pSI threshold = 1e-4).

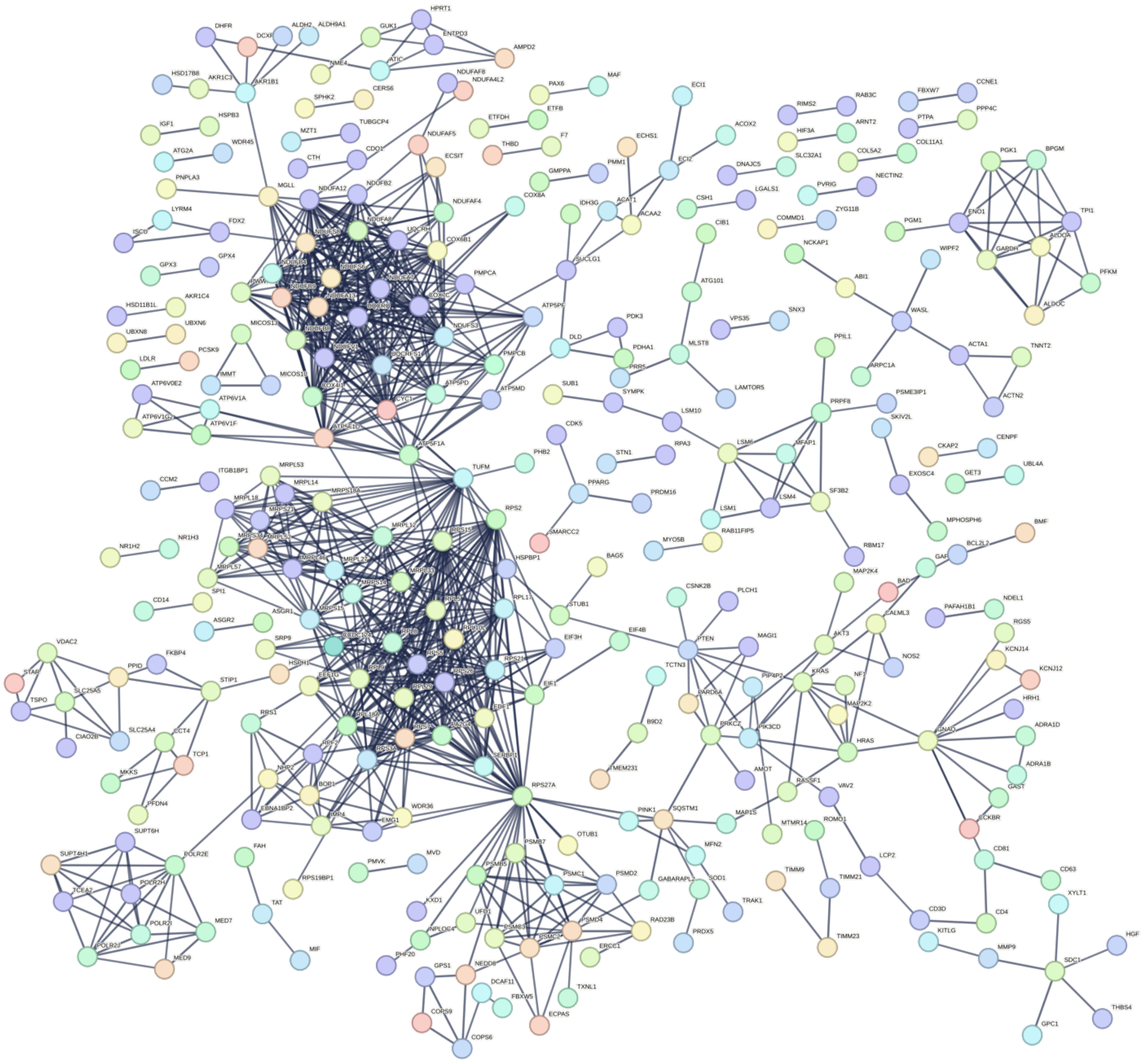

**Supplementary Figure 6.** A PPI network with statistical significance was constructed by the URGs related to cortical complexity changes in ASD.

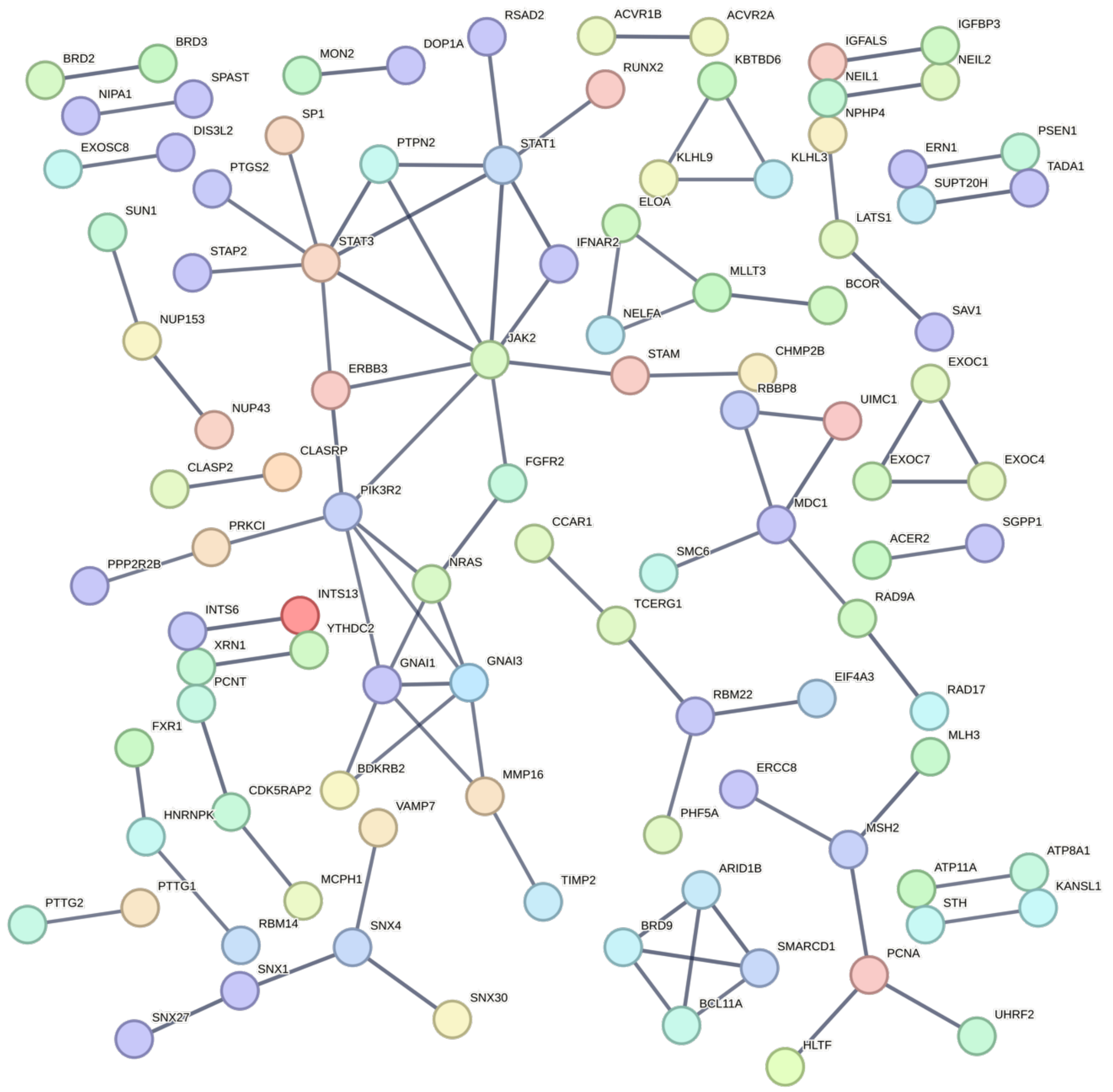


**Supplementary Figure 7.** A PPI network with statistical significance was constructed by the DRGs related to cortical complexity changes in ASD.

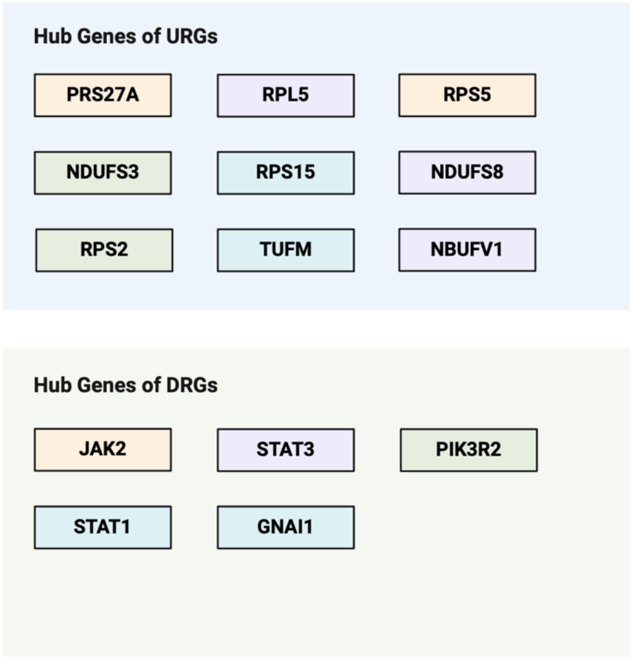


**Supplementary Figure 8.** Identified hub genes related to cortical complexity changes in ASD.
